# ARID1A and ARID1B preserve B cell identity, prevent myeloid transformation and reveal therapeutic vulnerabilities

**DOI:** 10.1101/2025.09.16.676393

**Authors:** Phyo Nay Lin, Joohyung Park, Yoon-A Kang, George P. Souroullas

**Affiliations:** Department of Medicine, Division of Oncology, Washington University School of Medicine, St. Louis, Missouri, USA; Department of Pediatrics, Washington University School of Medicine, St. Louis, Missouri, USA; Department of Medicine, Division of Hematology, Washington University School of Medicine, St. Louis, Missouri, USA; Siteman Comprehensive Cancer Center, Washington University School of Medicine, St. Louis, Missouri, USA

## Abstract

Chromatin remodeling by the SWI/SNF complex is essential for hematopoietic lineage commitment and differentiation. While core subunits *ARID1A* and *ARID1B* are frequently mutated in B cell malignancies, their functions remain unclear. Recent work established ARID1A- dependent functions within germinal center (GC) B cells, but its role during early B cell development, and whether its homolog, ARID1B, contributes distinct or compensatory roles at steady state or during transformation, remain unknown. Here, we used CD19-Cre-mediated deletion initiated at the pro-B cell stage to investigate their role in B cell development *in vivo*. Loss of either gene partially blocked B cell differentiation, reducing immature/recirculating B cell output, and impaired germinal center formation following antigen challenge. Combined deletion further reduced peripheral B cells, shortened survival, and resulted in aggressive leukemia. Unexpectedly, the malignancy was of myeloid origin and arose from a subset of CD19-expressing multipotent progenitors (MPPs). *Arid1a*/*Arid1b*-deficient MPPs exhibited abnormal expansion, reduced colony formation, and dysregulation of stemness and lineage-priming programs, including diminished CBFA2T3 (ETO2) and Fli1 signatures. In established B cell lymphoma cells *in vitro*, double ARID1A/ARID1B loss modestly affected cell growth, whereas loss of ARID1A increased sensitivity to EZH2 inhibition. Transcriptomic analyses revealed alterations in cell adhesion/migration pathways, cytokine-receptor interactions and DNA repair mechanisms. Collectively, these findings reveal stage-specific and compensatory roles for ARID1A and ARID1B in B cell development, uncover a mechanism by which SWI/SNF loss in MPPs redirects transformation towards myeloid leukemia, and suggest context-dependent therapeutic vulnerabilities.

## Introduction

The SWI/SNF complex, also known as BAF, is an ATP-dependent chromatin remodeling complex that modulates nucleosome positioning and chromatin accessibility. SWI/SNF is comprised of multiple subunits, with ARID1A and ARID1B serving as mutually exclusive core components of the canonical BAF complex. Across cancer, several BAF components function as tumor suppressors, especially in solid-tumors [1–3]. Mutations in different BAF subunits varies by cell type and tissue, implying subunit- and lineage-specific roles that also extend to hematopoietic malignancies of both myeloid and lymphoid origin.

Epigenetic and chromatin-modifying genes are essential regulators of hematopoietic lineage commitment and differentiation. ARID1A and ARID1B are highly expressed in hematopoietic stem and progenitor cells (HSPCs) and are commonly mutated in malignancies of B cell origin, with a combined mutation rate of about 20%, compared to about 2-3% mutation rate in myelodysplastic syndromes or acute myeloid leukemia (**Supp. Fig. 1**). How ARID1A and ARID1B contribute to hematopoietic transformation across lineages remains incompletely defined. In mice, *Arid1a* deletion in hematopoietic stem cells (HSCs) impairs stem cell frequency and multilineage differentiation, including B cell output [4]. However, because these deletions were initiated in HSCs, it remains unclear whether the B cell phenotype reflects intrinsic defects in B cell progenitors, or upstream perturbations in earlier progenitors. Loss of *Arid1b* in hematopoietic stem cells had a milder phenotype on steady-state hematopoiesis and HSC function [5,6]. Deletion of *Arid1a* or *Arid1b* was not associated with any malignancies in these studies. In mature B cells in the germinal center (GC) B cells, ARID1A-dependent canonical BAF orchestrates B cell fate during the GC response [7], is required for maintenance of GCs and high affinity antibody responses, and suppresses inflammatory gene programs [8]

Together, these observations point to stage-specific, context-dependent roles of the BAF complex in hematopoiesis and B cell neoplasia. However, several gaps remain in understanding its role during B cell development. For example, the role of BAF (with ARID1A or ARID1B) during B cell lineage commitment and B cell progenitor differentiation, remains unexplored. Furthermore, whether ARID1B provides compensatory or distinct functions during B-cell development, and how combined ARID1A/ARID1B loss affects lineage stability and malignant progression, has also not been addressed.

There is pressing need for new strategies to treat relapsed or refractory B cell malignancies. Epigenetic dysregulation is a hallmark of B cell neoplasms [9–17] and the reversibility of the epigenetic states presents promising therapeutic opportunities. Furthermore, emerging data also suggest that mutations in SWI/SNF components create therapeutic vulnerabilities, including synthetic lethal interactions with pharmacological inhibition of EZH2, CDK4/6 inhibition, and altered responses to immune checkpoint blockade [2]. These observations promoted the development of targeted therapies in clinical trials for SWI/SNF-deficient tumors [18–20], however, patient response is heterogeneous. Dissecting the lineage- and context-specific roles of ARID1A and ARID1B in hematopoietic cells may also inform future therapeutic strategies for B cell malignancies.

Extending beyond GC-restricted, ARID1A-only models, we used genetically engineered mouse models to delete *Arid1a* and *Arid1b* in B cell progenitors using the CD19-Cre, and assessed their role on B cell development, differentiation, germinal center (GC) responses, and potential role during malignant transformation. In parallel, we employed CRISPR/Cas9-edited human Diffuse Large B Cell Lymphoma (DLBCL) cells to test how double ARID1A/ARID1B loss affects growth of established lymphoma cells, shapes transcriptional programs, and modulates sensitivity to EZH2 inhibition. These complementary *in vivo* and *in vitro* approaches uncover both redundant and non-redundant roles of ARID1A and ARID1B in preserving B cell identity, reveal how early progenitor perturbations can redirect transformation towards myeloid leukemia, and identify context-dependent therapeutic vulnerabilities relevant to ARID1-mutant lymphomas.

## Materials and Methods

### Mice

All mice were backcrossed to the C57Bl/6 background and housed in an Association for Assessment and Accreditation of Laboratory Animal Care (AAALAC)-accredited facility and treated in accordance with protocols approved by the Institutional Animal Care and Use Committee (IACUC) for animal research at Washington University in St. Louis. Both male and female mice were included in all experiments. Tail genomic DNA was used for genotyping by PCR.

### Flow cytometry

Peripheral blood was collected into tubes containing potassium EDTA and complete blood counts were counted on a Hemavet (Drew Scientific). Flow cytometry analysis was carried out on Attune (Thermo) or FACS Aria (BD), using antibodies listed in Supp. Table 5. Data was analyzed using FlowJo software (TreeStar Inc.).

### Antigen response and germinal center analysis

For response to antigen, mice were injected with a solution of 2% sheep red blood cells (SRBCs) (Co-Caligo Biologicals) in 500μl PBC intraperitoneally (IP). Spleens were harvested and a single-cell suspension was analyzed by flow cytometry 9 days later. GC B cells were identified by expression of B220, GL7 and FAS.

### Bone marrow transplantation assays

Whole bone marrow was collected from femora and tibiae of donor CD19-Cre *Arid1a*^*F*/F^ *Arid1b*^*F*/F^ mice. Recipient mice were sub-lethally irradiated with a dose of 5Gy, and 5×10^5^ donor cells (CD45.2) were injected into each recipient mouse (CD45.1).

### Histology

Tissue samples were fixed in 10% formalin, processed for paraffin embedding by Histowiz, sectioned at 5 μm and stained with hematoxylin and eosin (H&E), and immunohistochemistry for PNA and Ki67.

### RNA sequencing and analysis

Appropriate cell populations were sorted from bone marrow using fluorescence-activated cell sorting (FACS) and processed for RNA extraction using Qiagen RNeasy Micro kit (Qiagen 74004). RNA-seq libraries were prepared following the manufacturer’s protocol, indexed, pooled, and sequenced on an Illumina NovaSeq 6000. Reads were base-called and demultiplexed using bcl2fastq2, and aligned to the Ensembl release 101 primary genome assembly. Gene counts were processed in R using the edgeR package, with TMM normalization applied to adjust for library size differences. Genes with low expression (defined as not expressed above one count-per-million in at least n–1 samples of the smallest group) and ribosomal genes were excluded. Normalized counts were transformed using voomWithQualityWeights in limma, incorporating surrogate variable analysis (SVA) to model latent sources of variation. Differential expression was assessed using linear modeling in limma, and results were filtered for FDR-adjusted p-values ≤ 0.05. Gene set enrichment analysis was performed using GAGE to identify perturbed GO, MSigDb, and KEGG pathways. Key gene modules were identified via weighted gene correlation network analysis (WGCNA) using limma-derived voom log2 counts-per-million values. RNA-seq data are available in Gene Expression Omnibus (GEO), GSE307527.

### Cell culture and generation of ARID1A/1B knock out SUDHL6 cell lines

Cells were grown in RPMI 1640 (Gibco) with 10% FBS (Millipore Sigma) and 1% penicillin-streptomycin (Genesee Scientific). Diffused large B cell lymphoma cells (SUDHL6) were transduced in 24-well plates with the TLCV2 lentivirus expressing target sgRNA guides along with inducible Cas9 and GFP. After 24 hours, transduced SUDHL6 cells were selected by exposure to 3µg/mL of puromycin for 48 hours. After selection, Cas9-2A-eGFP expression was induced by 1µg/mL doxycycline and GFP positive cells were sorted to establish clones. Knock out was validated by immunoblotting, qPCR and Sanger Sequencing.

### *In vitro* assays

For myeloid colony-forming assays, MPP populations were isolated by FACS (Aria, BD) and plated into M3231 methylcellulose media (Stem Cell technologies) supplemented with 20% IMDM media containing SCF (25 ng/ml), Flt3L (25 ng/ml), IL-11 (25 ng/ml), IL-3 (10 ng/ml), GM-CSF (10 ng/ml), EPO (4 U/ml), TPO (25 ng/ml) and 1% Pen/Strep. Colonies were counted and scored 7 or 8 days later. For Cell Migration Assays, Millicell Cell Culture Inserts 8.0 µm (Millipore Cat# PI8P01250) were used in 24-well plates in triplicate. 100 µL of serum-free medium was added into the upper chamber of the Millicell Cell Culture Inserts to hydrate the membrane, while 500,000 cells were suspended in 200 µL serum-free medium in the upper chamber and 750µL of culture medium with 20% FBS is placed in the lower chamber. The plate was incubated at 37°C for 24 hours. After 24 hours, the lower chamber medium was collected and analyzed by flow cytometry on an Attune analyzer (ThermoFisher). For cell growth assessment, the Alamar Blue assay was used. In rounded-bottom 96 well-plates, *ARID1A* and *ARID1B* knockout SUDHL-6 cells were seeded at 100 cells/well. Each condition was triplicated, and a non-target cell line is used as a control. Tazemetostat was used at 0nM, 100nM, 1µM, 5µM, and 10µM concentrations and cell growth was measured using the Alamar Blue assay at days 0, 5 and 10. Plates were analyzed on a SpectraMaxi3 instrument with fluorescence band width of 9 nm for excitation and 15 nm for emission.

### Western Blotting

DLBCL cells (2–3 × 10^6) were lysed on ice in 400 µL 1× RIPA buffer (CST #9806) with protease inhibitor (Thermo #A32953), sonicated, and clarified (14,000 g, 10 min, 4 °C). Protein was quantified by DC Protein Assay (Bio-Rad #5000112) using BSA standards, mixed 3:1 with 4× Laemmli buffer (Bio-Rad #1610747) + β-mercaptoethanol, and heated 10 min. Equal protein (200 µg/lane) was resolved on Mini-PROTEAN TGX gels (Bio-Rad #4561094; 100 V, ∼2 h) and transferred to PVDF in Tris-glycine-10% methanol (90 V, ∼2 h). Membranes were blocked (5% milk/TBS-T, 1 h), incubated with primary antibodies (1:1,000, 4 °C overnight), washed (TBS-T, 3×5 min), incubated with LI-COR secondary antibodies (IRDye #926-32211/#926-68070; 1:15,000, 1 h), washed, and imaged on a LI-COR Odyssey.

### Statistical analysis

Statistical analysis was performed using GraphPad Prism. Bar graphs represent mean and either standard deviation or standard error of the mean as indicated in the figure legends. Sample sizes were determined based on pilot studies or historical data from similar experiments. Power analysis was performed for two-tailed analysis, under the assumption of normal distribution, with a significance level of 0.05 and power of 0.8. Differences between groups were determined using Student’s t-test unless otherwise indicated.

## Results

### Deletion of *Arid1a* and *Arid1b* impairs B cells differentiation in the bone marrow

To investigate the role of *Arid1a* and *Arid1b* during B cell development we generated conditional knockout mice using the CD19-Cre allele, which is active from the pro B cell stage onward. Flow cytometric analyses of bone marrow and peripheral blood across multiple time points (4 weeks to ∼1 year of age) revealed a partial block in B cell differentiation in *Arid1a* and *Arid1b* single knockouts, with more pronounced effects in the double knockouts. Specifically, analysis of B cell progenitors in the bone marrow revealed that loss of either *Arid1a* or *Arid1b* resulted in an accumulation of pro B and pre B cells, and a reduction in immature and recirculating B cells (B220+, IgM^high^) (**Fig. 1a,b**), suggesting defects in differentiation or trafficking. Interestingly, peripheral blood B cell numbers remained largely unaffected in single knockouts, whereas double Arid1a/1b deletion caused a significant reduction in circulating B cells, and a drop in total white blood cell counts (**Fig. 1c-d**), indicating potential functional redundancy between *Arid1a* and *Arid1b* in the B cell lineage.

**Figure 1.**
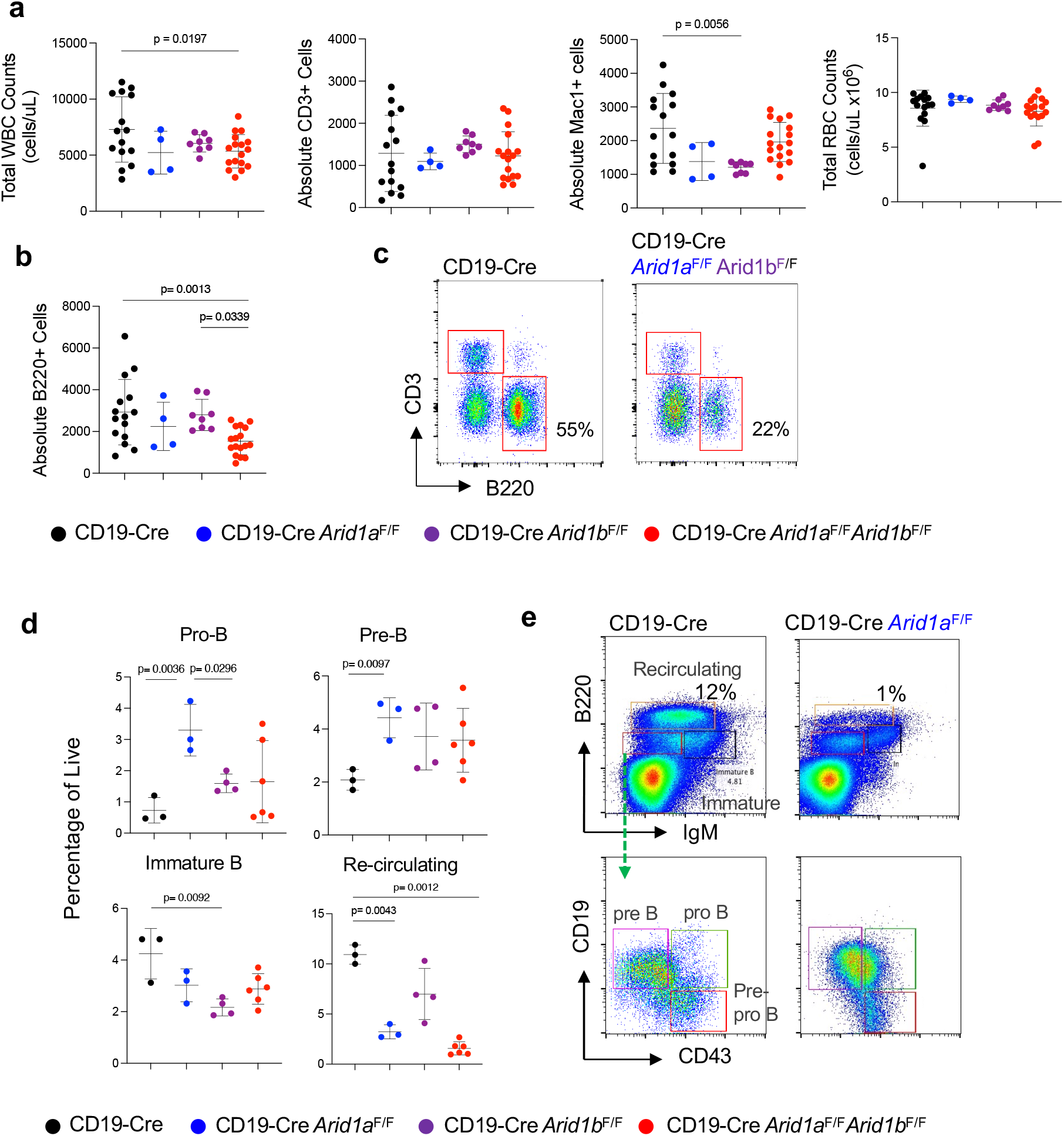
Deletion of *Arid1a* or *Arid1b* partially blocks B cell development and disrupts B cell output. **(a)** Peripheral blood analysis by flow cytometry and complete blood counts in the indicated genotypes. WBC=white blood cells, CD3+ T cells, Mac1+ myeloid cells, RBC=red blood cells. (n=5-15) **(b)** Absolute B cell counts in the indicated genotypes. (n=5-15) **(c)** Representative flow cytometric gating strategy used for panel (b) **(d)** Quantification of pro B, pre B, immature and recirculating B cell populations in the bone marrow of mice with the indicated genotype. (n=3-5). (**e**). Representative flow cytometry plots for panel (d). P values as indicated.

### *Arid1a* and *Arid1b* are required for Germinal Center expansion

Even though deletion of *Arid1a* or *Arid1b* did not have a measurable effect on B cell sub- populations in the spleen of young mice (data not shown), we hypothesized that Arid1a and Arid1b may play context-dependent roles in immune activation. To test this, we immunized 8-week-old mice with sheep red blood cells (SRBC) and assessed germinal center (GC) formation nine days later. Flow cytometry analysis of splenic B cells showed a significant reduction in Fas1+ GL7+ GC B cells in both CD19-Cre *Arid1a*^F/F^ and CD19-Cre *Arid1b*^F/F^ knockouts compared to CD19- Cre only controls (**Fig. 2a**). GC area as a fraction of total spleen was also significantly reduced (**Fig. 2b**). Immunohistochemistry showed a decrease in PNA+ cells, while Ki67 staining was similar among present cells, suggesting that the defect was not due to impaired proliferation but rather deficient GC formation or expansion.

**Figure 2.**
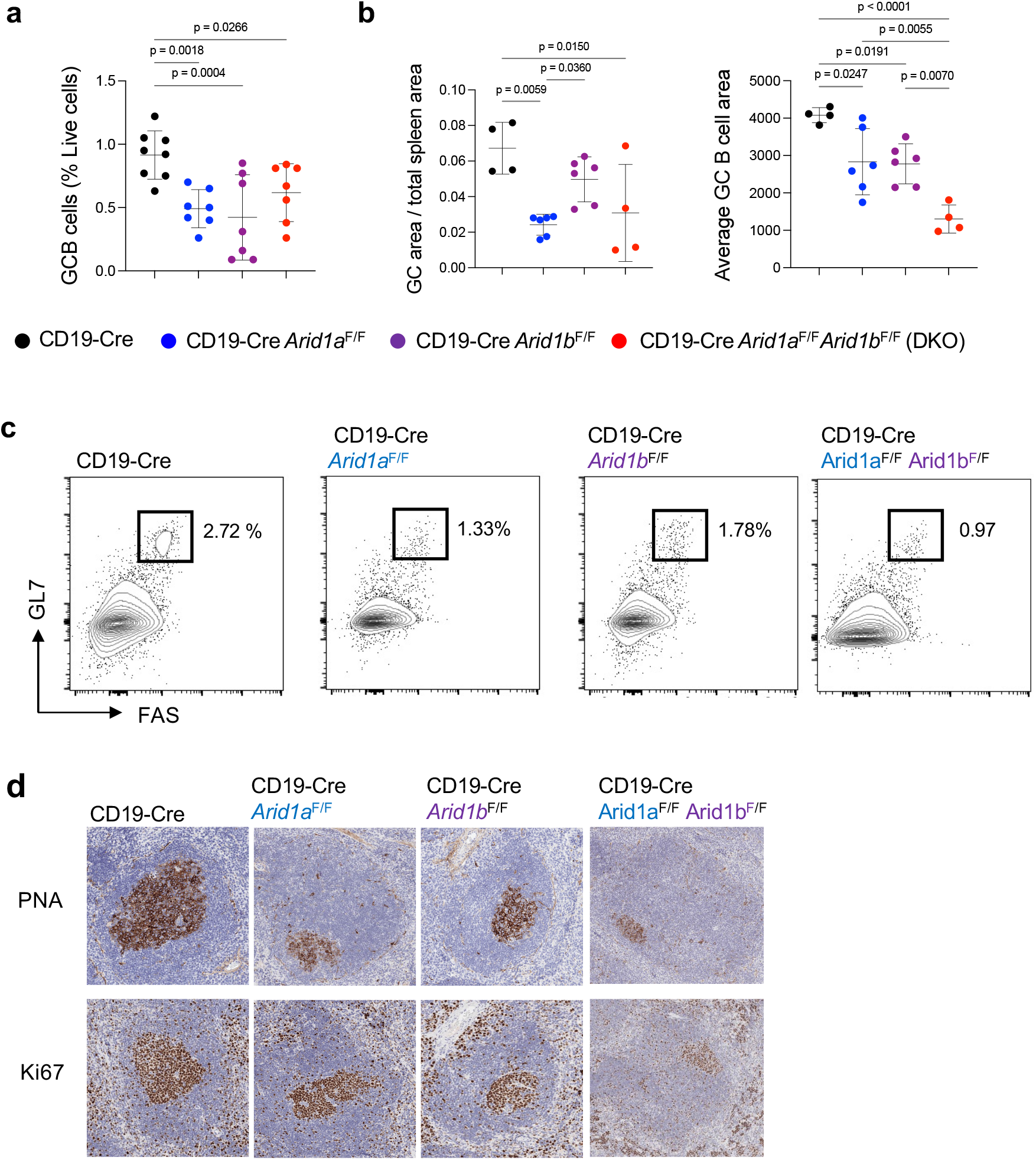
Deletion of *Arid1a* or *Arid1b* prevents germinal center expansion upon antigen exposure. **(a)** Quantification of GC B cells in the indicated genotypes as a percent of live cells in the spleen. n=7-8 **(b)** Calculation of GC area over the total spleen area (left), and the average GC area (right) (n=4- 6). **(c)** Representative flow cytometry of splenic B cells (B220+ Fas1+ GL7+) nine days after sheep red blood cell (SRBC) immunization from panel (a). **(d)** Immunohistochemistry for peanut agglutinin (PNA) and Ki67 in splenic germinal centers of *Arid1a*^*F*/F^, *Arid1b*^*F*/F^, and *Arid1a*^*F*/F^ *Arid1b*^*F*/F^ CD19-Cre only control mice

### Combined deletion of *Arid1a* and *Arid1b* using the CD19-Cre results in leukemia of myeloid origin

To assess the effect of *Arid1a* and *Arid1b* loss over long-term hematopoiesis, we monitored mice over two years. We found that CD19-Cre-mediated deletion of *Arid1a* and *Arid1b* resulted in significantly reduced survival, with double knockouts showing increased mortality by 3 months, and single knockout mice by 9 months of age (**Fig. 3a**, p<0.0001). These animals exhibited splenomegaly (**Fig. 3b**), leukocytosis, anemia, and thrombocytopenia. Since deletion of the *Arid1a/Arid1b* floxed alleles is driven by the CD19-Cre, which is expressed beginning at the pro B cell stage, we expected that these mice developed a type of B cell malignancy. Unexpectedly, flow cytometry revealed that the expanding population lacked expression of B cell markers (CD19-, B220-, IgM-, IgD-) and instead expressed markers of immature and myeloid cells (c-Kit+, CD43+, Mac1+, Gr1+) (**Fig. 3c-e**), suggesting transformation of a non-B lineage. Histological and immunophenotypic analysis of spleens showed high Ki67 expression and aberrant c-Kit^high^ populations (**Fig. 3d, Supp. Fig. 2**). To assess self-renewal capacity of this population, we transplanted bone marrow from diseased CD19-Cre *Arid1a*^F/F^*Arid1b*^F/F^ CD45.2 mice into sub lethally irradiated CD45.1 recipients. Donor-derived cells rapidly engrafted, maintained the aberrant immunophenotype, and led to recipient mortality (**Fig. 3e, f**), confirming the leukemic nature of the disease.

**Figure 3.**
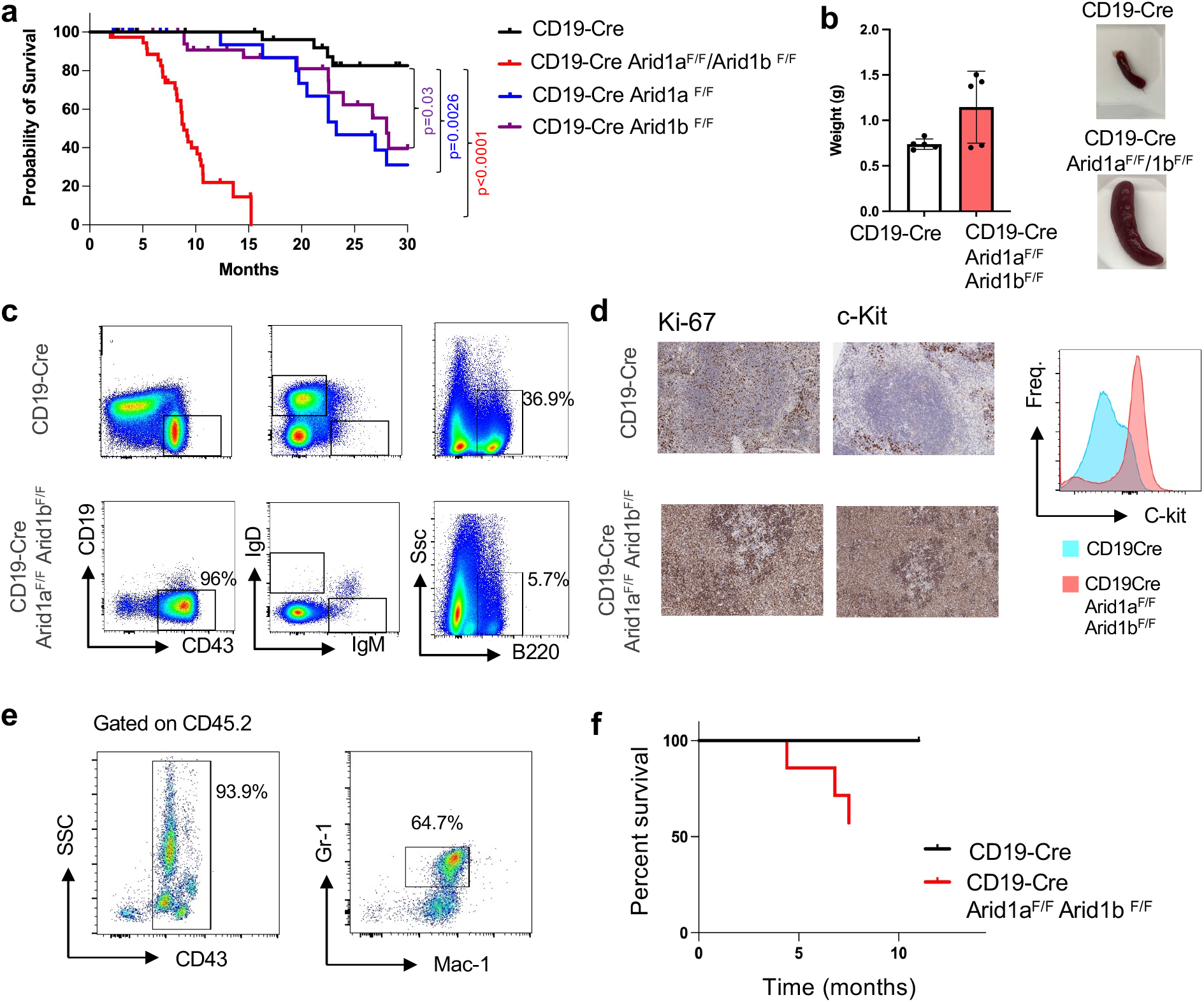
Deletion of both *Arid1a* and *Arid1b* results in shortened survival and myeloid malignancy. **(a)** Kaplan-Meier survival analysis of the indicated genotypes. Overall log-rank test p value<0.0001. P values of pair-wise log-rank tests of Arid1a/1b knockouts vs CD19-Cre control as indicated. (CD19-Cre n=35, CD19-Cre *Arid1a*^*F*/F^ n= 37, CD19-Cre *Arid1b*^*F*/F^ n=41, CD19-Cre *Arid1a*^*F*/F^ *Arid1b*^*F*/F^ n= 37) **(b)** Spleen weights and gross morphology of spleens demonstrating splenomegaly in moribund CD19-Cre *Arid1a*^*F*/*F*^*Arid1b*^*F*/F^ mice. n=5. **(c)** Representative flow cytometry analysis of spleens and bone marrow from CD19-Cre *Arid1a*^*F*/*F*^*Arid1b*^*F*/F^ mice for B cell markers (CD19, B220, IgM, IgD, CD43), highlighting expansion of abnormal immunophenotypic population. **(d)** Immunohistochemistry (IHC) for Ki67 and c-Kit in CD19-Cre *Arid1a*^*F*/*F*^*Arid1b*^*F*/F^ mice (left), and quantification of c-kit+ cells by flow cytometry (right). **(e)** Flow cytometry analysis of the blood of transplant recipient mice. **(f)** Kaplan-Meier survival analysis of transplant recipient mice. N=5

### Myeloid Leukemia arises from CD19-expressing multipotent progenitors

Given the myeloid phenotype of the developing leukemia, we hypothesized that the CD19-Cre maybe be active in a subset of multipotent progenitors (MPPs) with myeloid potential. To assess at what stage of hematopoietic differentiation CD19 may be expressed, we crossed the CD19- Cre to Lox-Stop-Lox TdTomato reporter mice. We found that ∼9% of MPP2 and ∼3% of MPP3 cells expressed TdTomato (**Fig. 4a-c**), and verified that sorted MPP2/3 populations had recombined the floxed *Arid1a* and *Arid1b* alleles (**Fig. 4d**).

**Figure 4.**
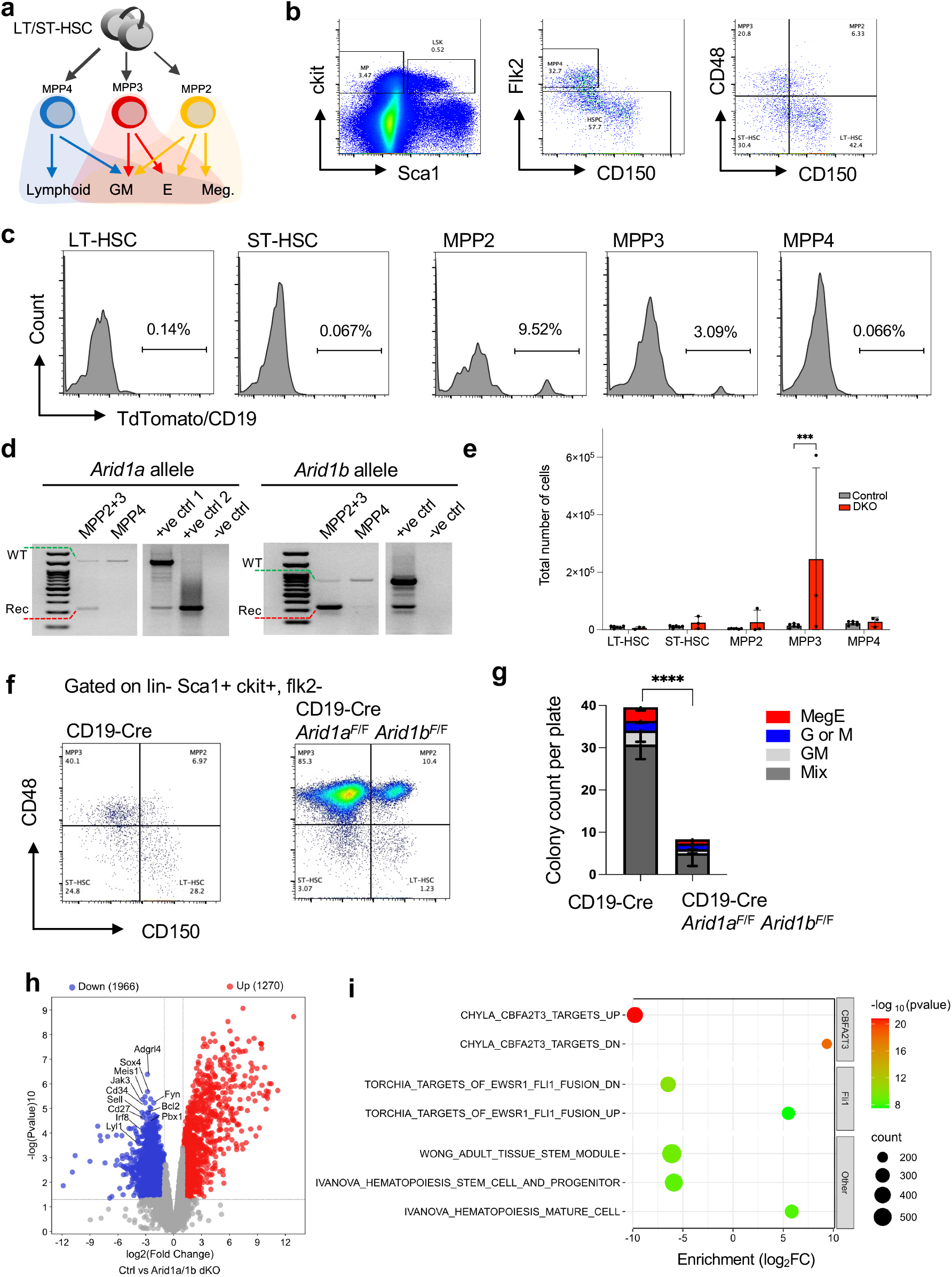
CD19-Cre-mediated deletion of *Arid1a* and *Arid1b* in multipotent progenitors drives expansion and differentiation defects. **(a)** Schematic of differentiation potential of multipotent progenitor (MPP) populations in the bone marrow. **(b)** Flow cytometric detection of HSCs and MPPs in CD19-Cre mice. **(c)** Representative flow cytometric detection of TdTomato reporter expression in CD19-Cre LSL-TdTomato mice in the indicated hematopoietic stem and progenitor populations from panel (**b**). n=3. **(d)** PCR genotyping of sorted MPP cells for recombination of the floxed *Arid1a* and *Arid1b* alleles in the indicated progenitor populations. WT= wild-type allele, Rec.= recombined allele. **(e)** Quantification of hematopoietic stem and progenitor cell populations in CD19-Cre and CD19- Cre *Arid1a*^*F*/*F*^*Arid1b*^*F*/F^ mice in 8-12month old mice. n=3-5. **(f)** Representative flow cytometry plots for panel (**e**). **(g)** Myeloid colony forming assays of MPP2/3 cells using methylcellulose. MegE, megakaryocyte or erythroid cells; G (or) M, granulocyte or macrophage; GM, granulocyte/macrophage; Mix; mix of all colonies (n=3-5), p<0.001. **(h)** Volcano plot showing differentially expressed genes in pooled MPP2/3 cells, comparing CD19- Cre vs CD19-Cre *Arid1a*^*F*/*F*^*Arid1b*^*F*/F^. **(i)** Gene expression signatures enriched or depleted in CD19-Cre *Arid1a*^*F*/*F*^*Arid1b*^*F*/F^ MPP2/3 cells compared to CD19-Cre controls, highlighting stemness and differentiation master regulators. LSL: lox-stop-lox, Td: Tandem dimer, lin: lineage cocktail (does not include CD19)

To determine how deletion of *Arid1a* or *Arid1b* affects MPP populations before the onset of leukemia, we analyzed the bone marrow of mice at different ages using flow cytometric and found progressive accumulation of MPP2/3 cells at the expense of HSCs and downstream progenitors (**Fig. 4e-f**). To determine if *Arid1a*/*Arid1b*-deficient MPP2/3 populations maintain their characteristic multipotency and robust colony-forming capacity, we isolated MPP2/3 and performed myeloid colony forming assays using methylcellulose as previously described [21]. Colonies were assessed at day 7 or 8. We found that *Arid1a/Arid1b*-deficient MPP2/3 cells formed significantly fewer colonies compared to controls, indicating impaired proliferation and/or differentiation capacity (**Fig. 4g**).

### Loss of *Arid1a* and *Arid1b* disrupts multipotency and differentiation programs in MPPs

To understand how genetic and molecular mechanisms change after *Arid1a*/*Arid1b* deletion in MPPs, we performed gene expression profile analysis via RNAseq on isolated MPP2/3 populations from CD19-Cre *Arid1a*^*F*/F^ *Arid1b*^*F*/F^ cells and CD19-Cre-only controls. Transcriptional profiling revealed downregulation of genes involved in stem cell maintenance and lineage priming (**Fig. 4h-i**), such as *Meis1, Pbx1, Lyl1* (**Fig. 4h**), and also significant downregulation of targets of *Cbfa2t3* (*Eto2*), and Fli1 (**Fig. 4i**), known regulators of stem cell fate decisions, and which are also directly implicated in the development of myeloid malignancies [22,23].

### ARID1A and ARID1B are dispensable for growth and proliferation in established B cell lymphoma cell lines

Although CD19-Cre deletion did not yield a B-cell malignancy *in vivo, ARID1A* and *ARID1B* are recurrently altered in mature B-cell lymphomas, more so than myeloid malignancies (**Supp. Fig. 1**). This suggests that they may shape disease maintenance, plasticity, or therapeutic response rather than initiation, a question best tested in an isogenic lymphoma context. To assess whether ARID1A or ARID1B are required for the proliferation of established B cell lymphoma cells, we used a lentiviral CRISPR/Cas9 system to generate targeted knockouts in the DLBCL cell line SUDHL-6 (**Supp. Fig. 3**). Single-guide RNAs targeting each gene were delivered alongside inducible Cas9 and GFP, and single-cell clones were isolated by FACs. For each gene, two independent sgRNAs were used to generate three clones per guide, and multiple double knockout (ARID1A/ARID1B) clones were also established. Control clones were derived using two non-targeting sgRNAs. Cell growth assays showed that deletion of *ARID1A* or *ARID1B* did not affect growth or proliferation compared to non-targeting controls. Combined deletion had a modest synergistic effect of cell growth (**Fig. 5a**), suggesting that they are not essential for cell-autonomous proliferation in this context.

**Figure 5.**
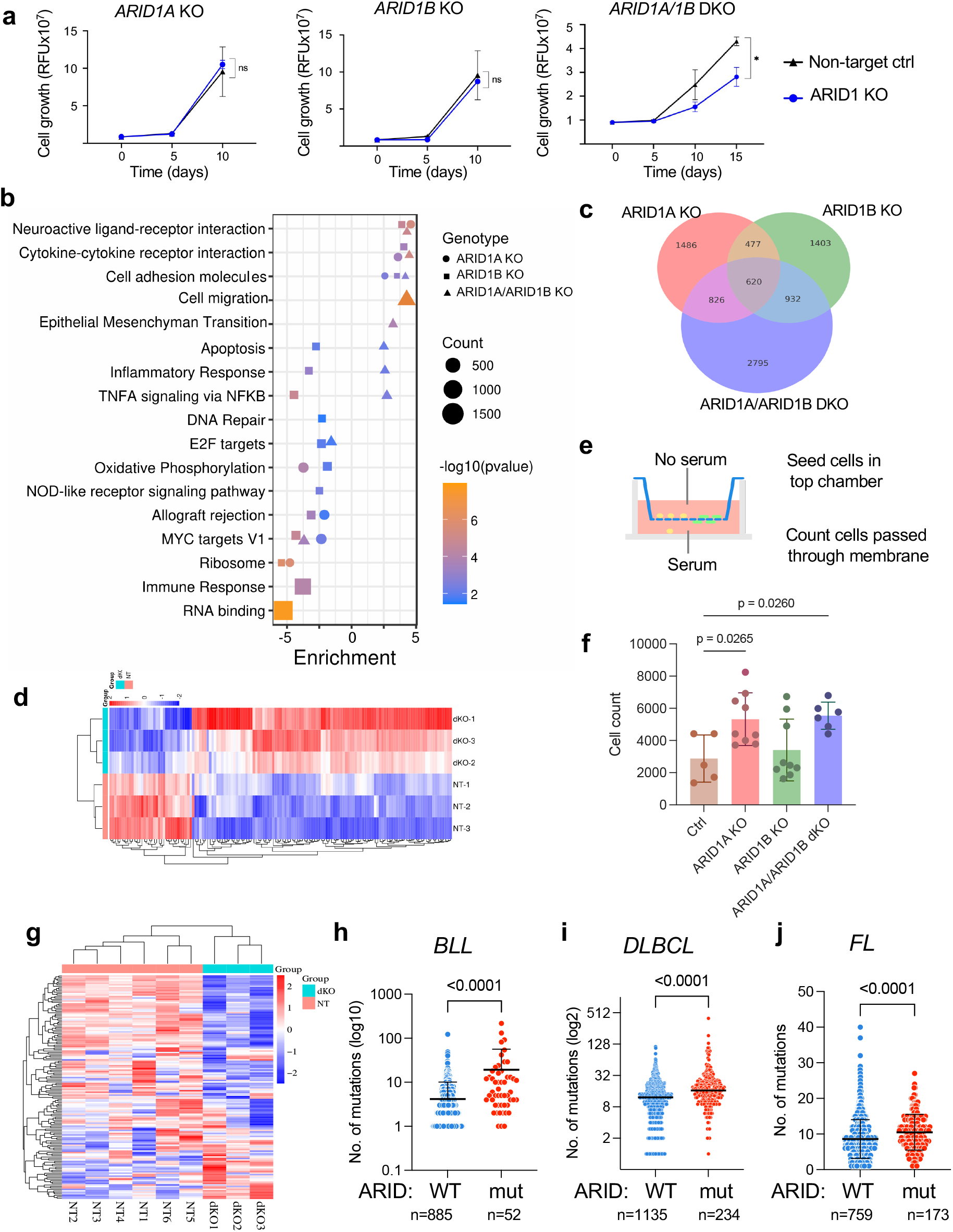
Deletion of *ARID1A* and *ARID1B* in established lymphoma cell line SUDHL6 enhances cell migration and is associated with increased mutation burden in patients. **(a)** *In vitro* growth curves of SUDHL6 cells after CRISPR/Cas9-targeting of *ARID1A* (left), *ARID1B* (middle) or both (right). KO=knockout **(b)** Pathway analysis (GAGE) of gene expression changes after *ARID1A, ARID1B* and double *ARID1A*/*ARID1B* deletion compared to non-target controls. **(c)** Overlapping gene expression changes in *ARID1A* KO, *ARID1B* KO and double *ARID1A*/*ARID1B* KO SUDHL6 cells (p<0.001, log_2_FC>2) **(d)** Heat map of cell migration signature in non-target control vs double ARID1A/ARID1B KO SUDHL6 cells. **(e)** Experimental schematic of *in vitro* transwell migration assay. **(f)** Quantification of transwell migration assays. **(g)** Heat map of the Hallmark DNA damage gene expression signature in non-target controls vs double ARID1A/ARID1B-targeted SUDHL6 cells. (**h-j**) Mutation burden analysis of patients that express either a wild-type vs mutated form of *ARID1A* or *ARID1B* in (h) B cell leukemia (BLL), (i) Diffuse large B cell lymphoma (DLBCL), or (j) Follicular Lymphoma (FL) patients. BLL: ARID1A/1B WT n=885, mut n=52. DLBCL: ARID1A/1B WT n=1135, mut n=234. FL: ARID1A/1B WT n=759, mut n=173. Mut: mutant.

### Loss of ARID1A or ARID1B alters expression of shared and distinct transcriptional programs

To investigate the effect of ARID1A and ARID1B deletion on molecular mechanisms in lymphoma cells, we performed RNA-sequencing. For each gene, two distinct sgRNAs were used to create three independent clones per guide (n=6 per gene), and non-targeting sgRNA clones were used as controls (n=6). Transcriptomic profiling revealed widespread changes in gene expression following deletion of each ARID1 homolog (**Supp. Fig. 4a**). While hundreds of genes were differentially expressed in each comparison (FDR < 0.05), there was substantial overlap between ARID1A and ARID1B knockouts (**Fig. 5b-c**). Pathway enrichment analysis using the Generally Applicable Gene-set Enrichment (GAGE) [24] method identified shared enrichment of pathways related to ribosome biogenesis, oxidative phosphorylation, E2F signaling, cell adhesion, cytokine-receptor interactions, DNA repair, and transcriptional networks regulated by MYC, EZH2 and other transcription factors (**Fig. 5b**). These findings suggest that both ARID1A and ARID1B contribute to global regulatory networks supporting proliferation and metabolic homeostasis in B cell lymphoma cells. Each gene also exhibited distinct transcriptional effects. *ARID1B* knockout clones showed enrichment for transcription factor signatures including IL2/STAT5 signaling, BMP2, and WT1, while ARID1A knockout clones were enriched for DREAM targets (**Supp. Fig. 4b**). These gene-specific differences may reflect functional specialization between ARID1A and ARID1B and underscore the context-dependent complexity of SWI/SNF regulation in lymphoma.

### Single deletion of *ARID1A* or double *ARID1A*/*ARID1B* deletion results in enhanced lymphoma cell migration *in vitro*

Several of the altered pathways identified in *ARID1A/ARID1B*-deficient lymphoma cells involved cell migration and adhesion mechanisms (**Fig. 5b, e**). While lymphomas are not typically considered solid tumors, cell migration and adhesion pathways still play crucial roles in disease biology such as cell trafficking, dissemination and interactions with the microenvironment [25]. To test whether deletion of *ARID1A* or *ARID1B* affect cell migration, we performed *in vitro* transwell migration assays. We found that deletion of *ARID1A* and double *ARID1A/ARID1B* resulted in increased cell migration (**Fig. 5f-g**), while deletion of *ARID1B* did not, suggesting that perhaps ARID1A/1B deletions contribute to disease pathogenesis through lymphoma dissemination.

### *ARID1B*-deletion alone is sufficient to cause alterations in DNA repair gene expression programs

Another molecular category that was particularly downregulated after *ARID1B* deletion was DNA repair mechanisms (**Fig. 5b**). We hypothesized that in lymphomas with *ARID1A* or *ARID1B* loss-of-function mutations, disrupted DNA repair mechanisms would result in increased mutation burden. Mutation rates *in vitro* may not be reliable because of the short-term nature of *in vitro* experiments after gene deletion relative to long-term disease progression in people. We thus assessed mutation burden from patients with B cell neoplasms. We analyzed patient sequencing data from The Cancer Genome Atlas (TCGA) and AACR Project GENIE v18.0 via cBioportal [26– 28], and stratified patients into two groups: *ARID1A*/*ARID1B* wild-type vs *ARID1A*/*ARID1B* mutants (truncating, missense or non-sense mutations), and assessed mutation burden. We found that in three different types of B cell neoplasms, (B cell lymphoblastic leukemia (BLL), Diffuse Large B cell Lymphoma (DLBCL) and (Follicular Lymphoma (FL), there was significantly higher mutation burden in patients with *ARID1A* or *ARID1B* mutation (**Fig. 5h-j**). These findings are consistent with previous reports implicating SWI/SNF components in DNA repair in solid tumors [29–31], either through physical interaction with the DNA damage response machinery (e.g., ATR) or by regulating expression of repair genes.

### Loss of ARID1A, but not ARID1B, sensitizes lymphoma cells to EZH2 inhibition

Finally, deletion of *ARID1A* and *ARID1B* resulted in depletion of nucleosome organization and chromatin assembly pathways, along with downregulation of some of its own components, such as SMARCC1 (**Fig. 6a-b**). We found significant upregulation of canonical PRC2 and EZH2 targets (**Fig. 6a**), and that EZH2 activity represses expression of SWI/SNF components such as SMARCC1, which we verified by western blotting (**Fig. 6b**). Consistent with a SWI/SNF-PRC2 antagonism model [32–35], multiple ongoing or completed trials have tested EZH2 inhibition in ARID1A-mutant or SWI/SNF-deficient tumors. In these trials, EZH2 inhibition showed some clinical activity, however, objective responses were limited [18–20], suggesting that EZH2- inhibitor sensitivity may not only be dependent on patient genetic/mutation diversity, but also SWI/SNF subunit composition. To test sensitivity to EZH2 inhibition in DLBCL cells, and to determine whether *ARID1A* or *ARID1B* have a similar or distinct outcomes after EZH2 inhibition, we treated *ARID1A*- and *ARID1B*-deficient SUDHL6 clones with the EZH2 inhibitor Tazemetostat. We found that deletion of *ARID1A*, and double *ARID1A/ARID1B* deletion rendered some clones more sensitive to EZH2 inhibition, while deletion of *ARID1B* alone did so to a lesser extent (**Fig. 6c-d**). These results highlight the fact that not only cellular context, but the composition of SWI/SNF itself may determine the crosstalk and synthetic lethality that exists between loss of SWI/SNF function and pharmacological inhibition of EZH2.

**Figure 6.**
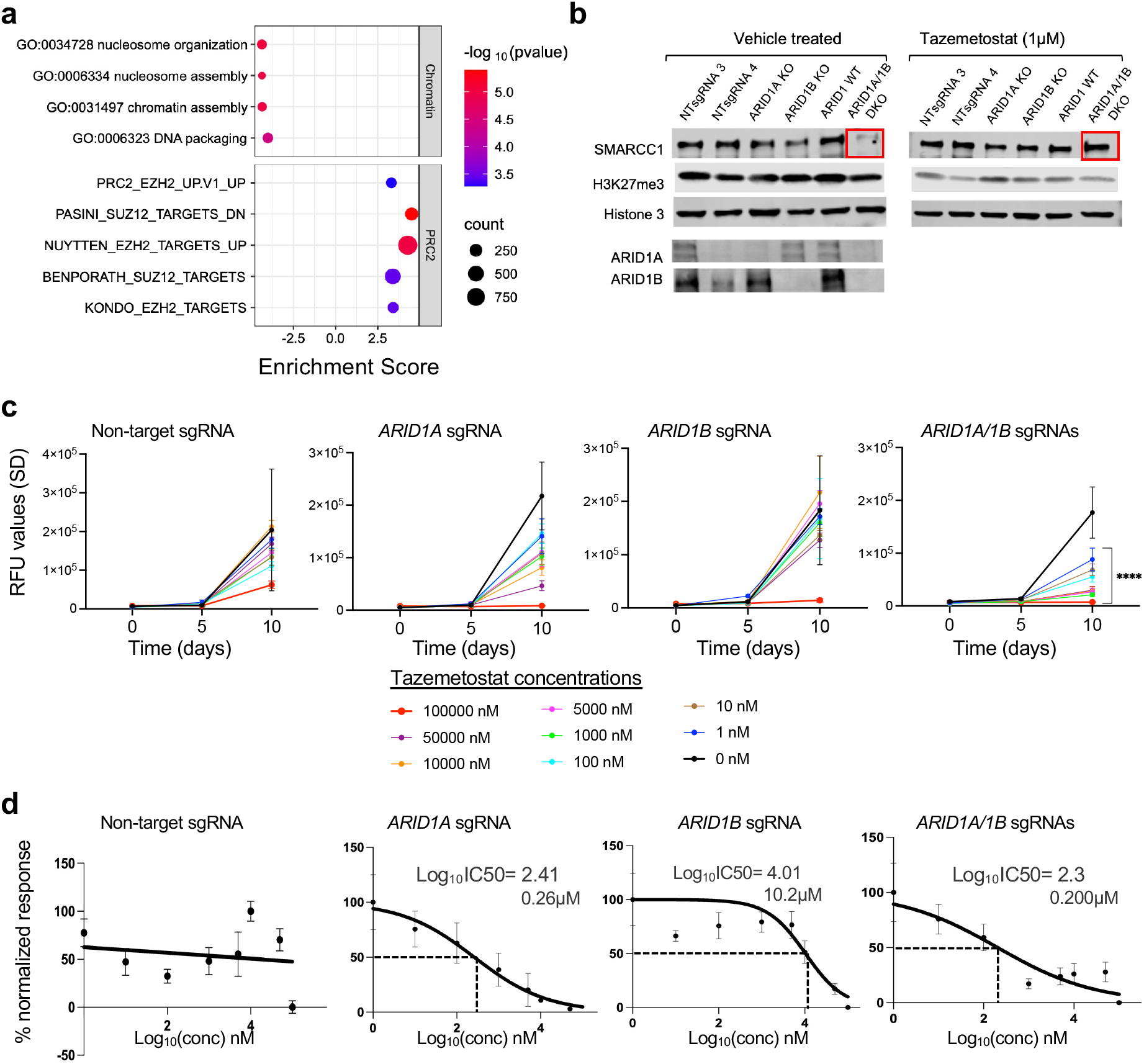
Effect of pharmacological EZH2 inhibition on double *ARID1A*/*ARID1B*-KO SUDHL6 cells. **(a)** Gene expression signatures in ARID1A/ARID1B double knockout SUDHL6 cells compared to non-target controls, highlighting loss of nucleosome organization pathways and enrichment of PRC2/EZH2 targets. **(b)** Western blot for indicated proteins from vehicle-treated and Tazemetostat-treated clones. (**c-d**) *In vitro* growth curves (c) and IC_50_ drug response curves (d) of non-target control vs ARID1A/1B targeted lines as indicated.

## Discussion

Our findings reveal critical and overlapping roles for the SWI/SNF subunits ARID1A and ARID1B in B cell development, germinal center function, and leukemogenesis. Using a lineage-specific conditional knockout model, we demonstrate that loss of either Arid1a or Arid1b in mice impairs early B cell differentiation in the bone marrow and compromises germinal center formation following antigen stimulation, while combined loss unmasks substantial redundancy, markedly reducing B cell output and shortening survival. Recent studies have shown that ARID1A- dependent canonical BAF activity is required to sustain germinal center (GC) programs and restrain inflammatory gene expression during the GC response [8], findings that align with our GC phenotypes for Arid1a loss, but do not address Arid1b or combined deficiency. These observations, together with our genetic interaction data, highlight both the dosage sensitivity and compensatory function of ARID1A and ARID1B in the B cell lineage.

Unexpectedly, combined deletion of *Arid1a* and *Arid1b* using the CD19-Cre resulted in a lethal leukemia of myeloid rather than lymphoid origin. This result prompted closer examination of CD19 expression during early hematopoiesis, revealing low-level Cre activity in multipotent progenitor populations (MPPs) that are sufficient to disrupt lineage fidelity and drive malignant transformation. Arid1a/Arid1b-deficient MPP2/3 subpopulations exhibited significant expansion, impaired differentiation, and transcriptional dysregulation of stem cell programs, including signatures associated with Eto2 (Cbf2t3) and Fli. While these alterations are correlative, they are consistent with prior roles of ETO2 and Fli1 in progenitor function and myeloid fate decisions, and together support a model in which SWI/SNF safeguards early lineage boundaries to prevent inappropriate myeloid conversion and transformation.

Our results also offer insight into the context-dependent tumor suppressor functions of ARID1A and ARID1B. Although frequent mutations in these genes have been documented in mature B cell lymphomas, our study indicates that their loss may also disrupt earlier developmental trajectories and prime progenitor cells for alternative fates. This dual role in lineage specification and transformation may explain why SWI/SNF mutations are observed across a spectrum of hematologic malignancies, including both lymphoid and myeloid subtypes. Moreover, our observation that leukemic transformation can arise from a progenitor population harboring early recombination events has implications for interpreting lineage-specific knockout models and for understanding disease heterogeneity in human cancers.

Our *in vitro* studies extend these concepts to established lymphoma cells. In DLBCL cells, loss of ARID1A or ARID1B reprogrammed overlapping transcriptional networks involved in ribosome biogenesis, oxidative phosphorylation, adhesion/migration, cytokine signaling, DNA repair, and EZH2-linked pathways. Phenotypically, ARID1A loss enhanced transwell migration, whereas ARID1B loss had minimal effect, suggesting that ARID1A may be the main regulator of migratory programs that could contribute to disease dissemination *in vivo*.

Therapeutically, our data support context-dependent antagonism between SWI/SNF and PRC2. Double ARID1A/ARID1B loss enriched PRC2/EZH2 target signatures and depleted nucleosome assembly programs. Functional assays showed increased sensitivity to EZH2 inhibition with ARID1A loss, whereas ARID1B alone had a smaller or variable effect. These findings suggest that SWI/SNF composition, not deficiency per se, modulates PRC2 dependency in lymphoma. This also implies that ARID1A might serve as a more informative biomarker of EZH2-inhibitor sensitivity than ARID1B alone

In summary, ARID1A and ARID1B cooperatively preserve B-cell identity and prevent myeloid transformation by stabilizing progenitor transcriptional programs. In established lymphoma their loss reshapes migration, DNA-repair, and PRC2-linked networks that reveal actionable dependencies. These insights argue for biomarker-guided stratification of EZH2-directed therapies by SWI/SNF subunit status and motivate combination approaches that integrate epigenetic and DNA-damage–response agents in ARID1-mutant B-cell malignancies.

## Acknowledgements

We thank the Siteman Cancer Center Flow Cytometry Core and the McDonnell Genome Institute/Genome Technology Access Center at Washington University for flow cytometry services and genome analysis respectively, and the Department of Comparative Medicine for animal maintenance and expertise. This project was funded in part by the Alvin J. Siteman Cancer Center, Barnes-Jewish Hospital and the Washington University Institute of Clinical and Translational Sciences, which is, in part supported by an NCATS Clinical and Translational Sciences Award, #UL1TR002345. The Siteman Cancer Center is supported in part by an NCI Cancer Center Support Grant #P30CA091842. Additional support was provided by the Barnard Cancer Institute.

## Authorship

Contributions: PNL, JP, YK and GPS performed experiments and analyzed data. PNL and GPS wrote the manuscript. GPS conceptualized and supervised the project.

