## Supplementary material for "ARID1A and ARID1B preserve B cell identity, prevent myeloid transformation and reveal therapeutic vulnerabilities": Supp. Figure legends

**Supplemental Figure Legends**

**Supp. Figure 1 . Mutation frequencies and clinical outcomes of patients with *ARID1A* or *ARID1B* alterations**

(**a**) OncoPrint of mutations in SWI/SNF components, mainly ARID1A, ARID1B and ARID2, in diffuse large B cell lymphoma (DLBCL), Myelodysplastic syndrome and Acute Myeloid Leukemia

(**b-c**). Ann-Arbor staging (b) and percentage of aggressive, stage IV cases of DLBCL patients with mutations in ARID1A, ARID1B, or both.

**(d)** Kaplan-Meier survival curve of DLBCL patients based on ARID1A/ARID1B mutation status. Source: Patient data from cBioportal analyzed for staging.

**Supp. Figure 2. Immunophenotyping of organs infiltrated with leukemia**

Immunohistochemistry for MPO, ckit, CD20 and Ki67 of spleen and liver section isolated from control CD19-Cre, CD19-Cre Arid1a$^{F/F}$ and CD19-Cre *Arid1a*$^{F/F}$*arid1b*$^{F/F}$ mice.

**Supp. Figure 3. CRISPR/Cas9-mediated targeting of ARID1A and ARID1B in SUDHL6 cells**

**(a)** ARID1A targeting was carried out using a doxycycline-inducible lentiviral/GFP CRISPR/Cas9 system. Image shows robust expression of Cas9-GFP

(**b**) Western blots verifying ARID1A or ARID1B deletion (NT=nontarget control).

(**c-d**) Sanger sequencing of the *ARID1A* (c) and *ARID1B* (d) loci, indicating location of the single guide RNA and the resulting mutation or deletion.

**Supp. Figure 4. Differential gene expression analysis of ARID1A-, ARID1B- and double ARID1A-1B-targeted SUDHL6 cells**

**(a)** Volcano plots of differential expression between non-targe control and *ARID1A*-targeted (left), *ARID1B*-targeted (middle) and double ARID1A/*ARID1B*-targeted (right) cells

(**b**) Unique enriched gene signatures after ARID1A (top) or ARID1B (bottom) deletion in SUDHL6 cells.
