## Supplementary material for "ARID1A and ARID1B preserve B cell identity, prevent myeloid transformation and reveal therapeutic vulnerabilities": Supp. Figures

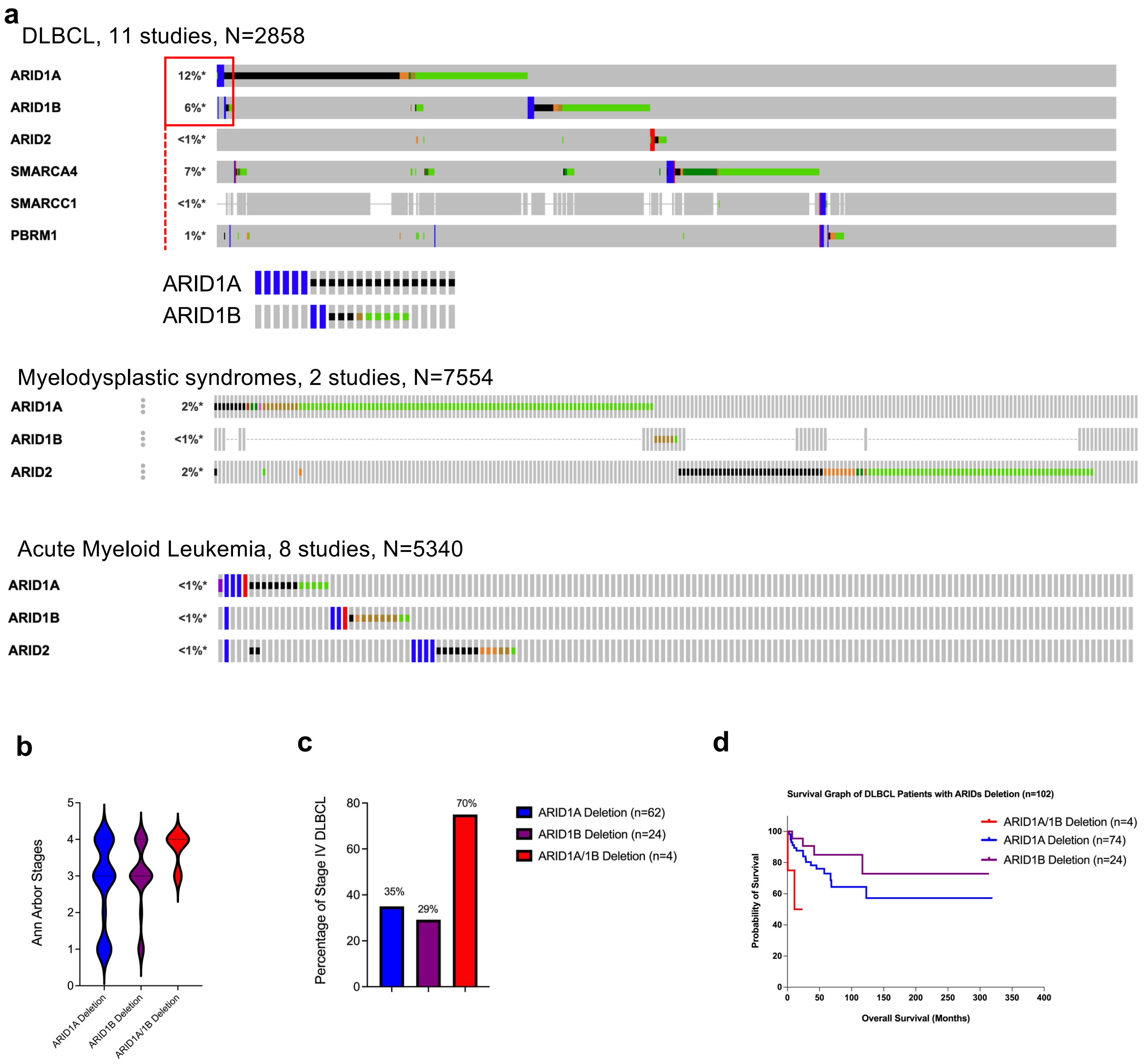


**Supp. Figure 1.** Mutation frequencies and clinical outcomes of patients with *ARID1A* or *ARID1B* alterations

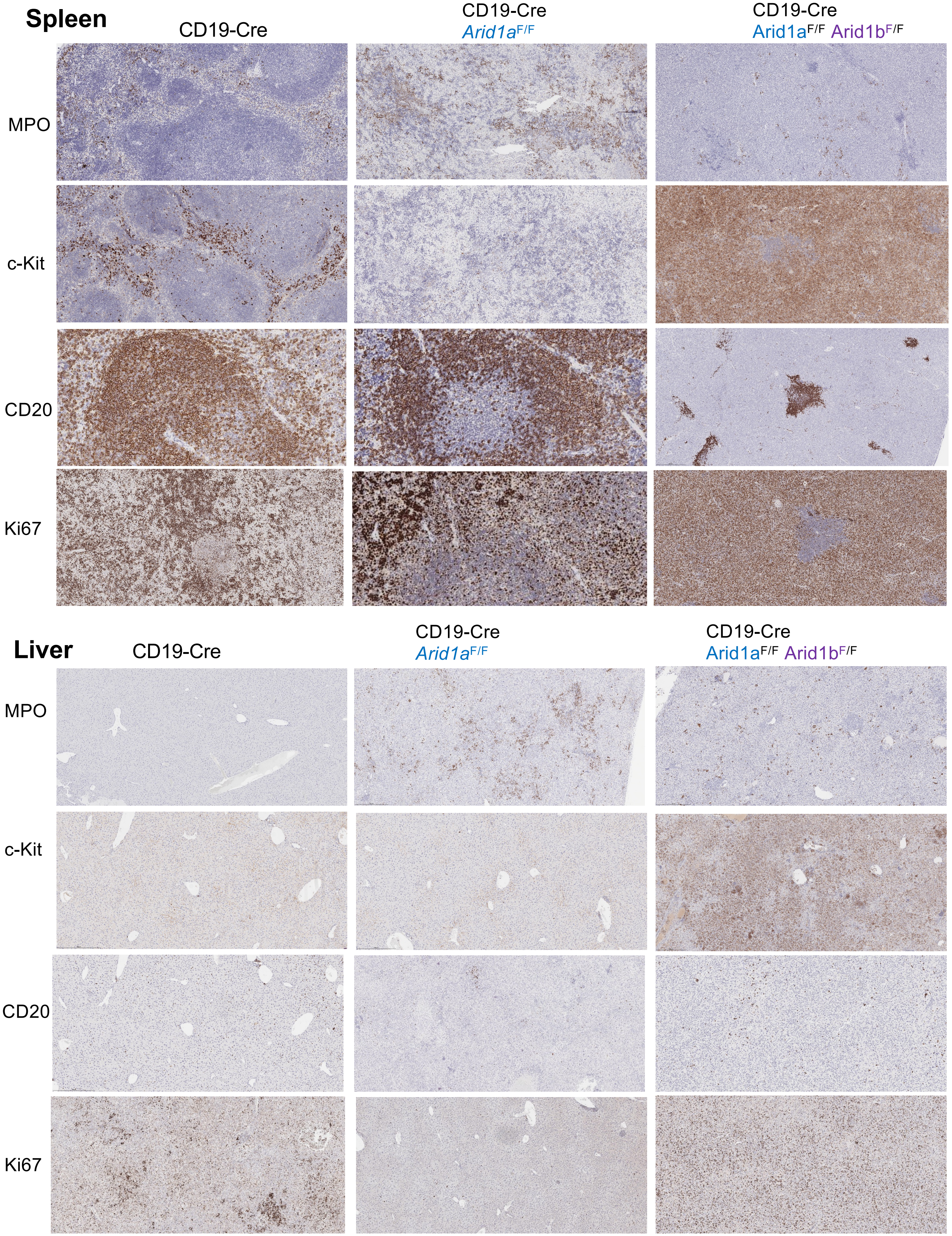


**Supp. Figure 2.** Immunophenotyping of organs infiltrated with leukemia

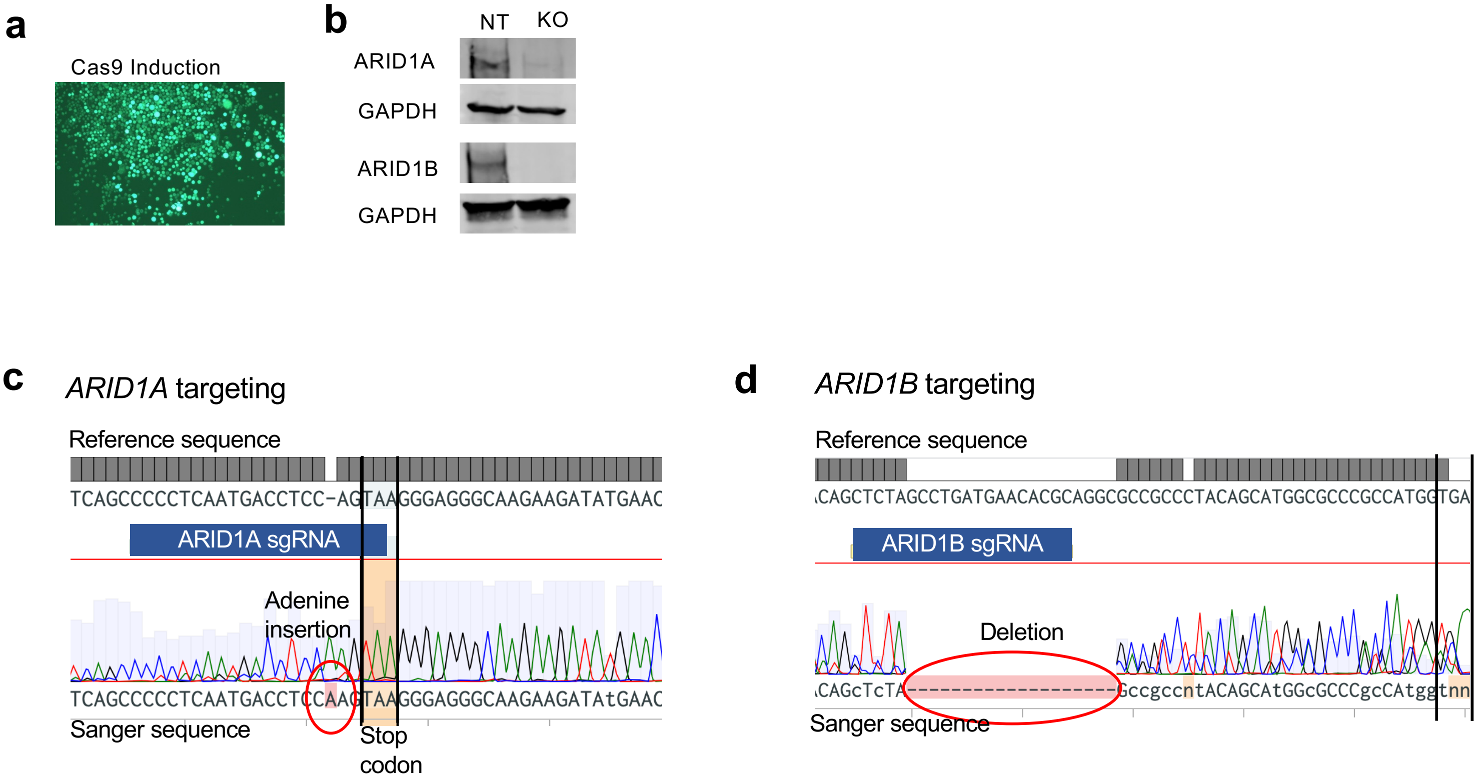


**Supp. Figure 3.** CRISPR/Cas9-mediated targeting of *ARID1A* and *ARID1B* in SUDHL6 cells

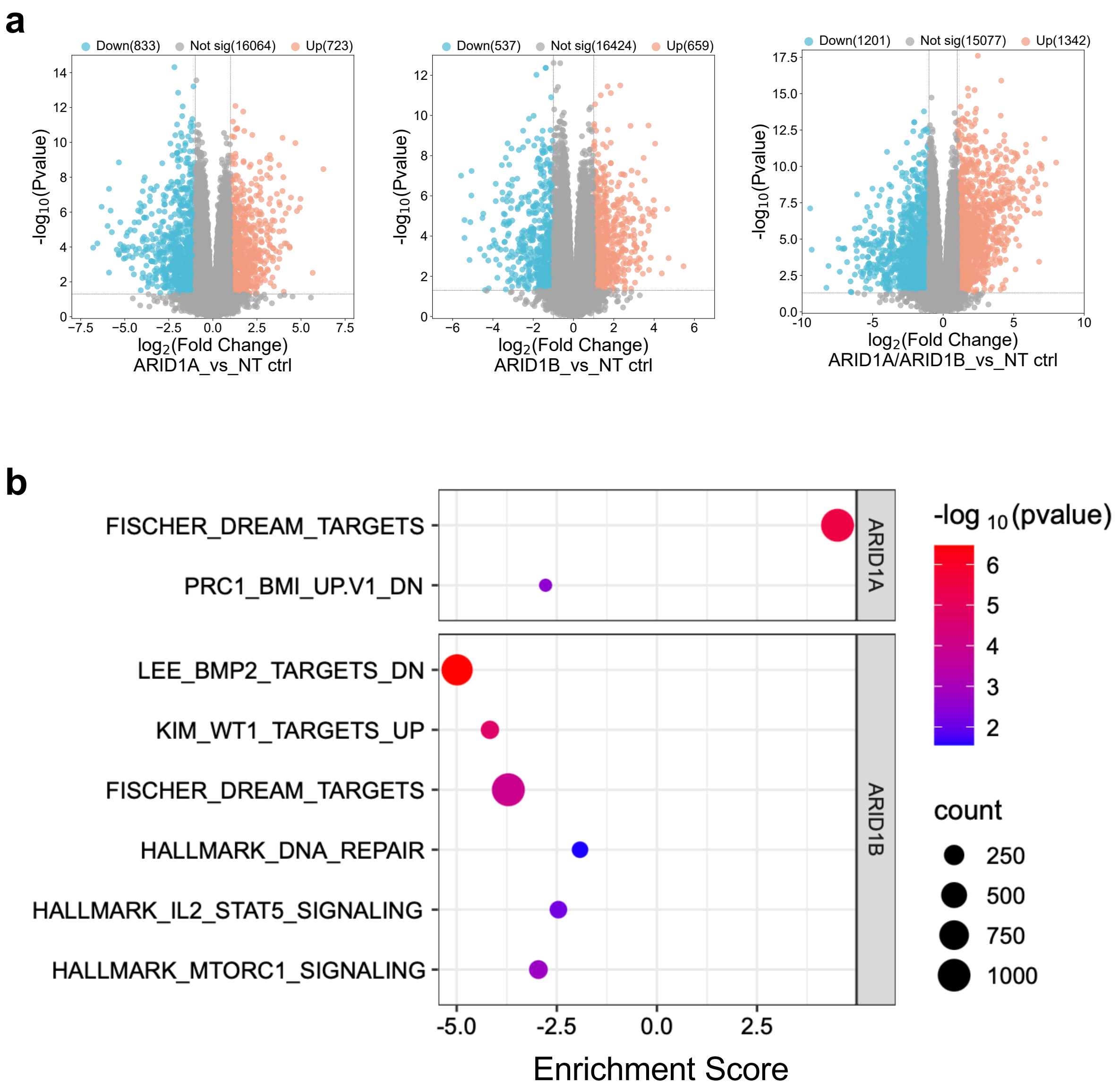


**Supp. Figure 4.** Differential gene expression analysis of ARID1A-, ARID1B- and double ARID1A-ARID1B-targeted SUDHL6 cells
