## Supplementary material for "ARID1A and ARID1B preserve B cell identity, prevent myeloid transformation and reveal therapeutic vulnerabilities": Tables

| Primers name | Sequence | Source |
|---|---|---|
| Cre MB182-F | CAATGGTAGGCTCACTCTGGGAGATGATA | IDT |
| Cre MB182-R | AACACACACTGGCAGGACTGGCTAGG | IDT |
| Cre MB183-F | CAGGGTGTTATAAGCAATCCC | IDT |
| Cre MB183-R | CCTGGAAAATGCTTCTGTCCG | IDT |
| Arid1a flox-F | CTAGGTGGAAGGTAGCTGACTGA | IDT |
| Arid1a flox-R | TACACGGAGTCAGGCTGAGC | IDT |
| Arid1b flox-F | CTTGGTCTTACCCATTTGCAC | IDT |
| Arid1b flox-R | ATCGATGGAGCCAGACAGGT | IDT |
| Arid1a Recombination-F | CTAGGTGGAAGGTAGCTGACTGA | IDT |
| Arid1a Recombination-R | AGAGTAACTAATAACTGCTGGAGGATG | IDT |
| Arid1b Recombination-F | CTTGGTCTTACCCATTTGCACAGT | IDT |
| Arid1b Recombination-R | GATGGAGGATCCTTACTACAGGGGGATT | IDT |

**Table 1. Genotyping primers**

| Name | Sequence | Source |
|---|---|---|
| Arid1a-sgRNA1-O1<br>Arid1a-sgRNA1-O2<br>Arid1a-sgRNA2-O1<br>Arid1a-sgRNA2-O2<br>Arid1a-sgRNA3-O1<br>Arid1a-sgRNA3-O2 | CACCGAAGAACTCGAACGGGAACGC<br>AAACGCGTTCCCGTTCGAGTTCTTC<br>CACCGGCGGTACCCCATGACCATGC<br>AAACGCATGGTCATGGGGTACCGCC<br>CACCGTTAGTCCCACCATACGGCTG<br>AAACCAGCCGTATGGTGGGACTAAC | IDT |
| Arid1b-sgRNA1-O1<br>Arid1b-sgRNA1-O2<br>Arid1b-sgRNA2-O1<br>Arid1b-sgRNA2-O2<br>Arid1b-sgRNA3-O1<br>Arid1b-sgRNA3-O2 | CACCGCGCGCAACAAAGGAGTCACC<br>AAACGGTGACTCCTTTGTTGCGCGC<br>CACCGCTGTGCACCTGGGGGACCGT<br>AAACACGGTCCCCCAGGTGCACAGC<br>CACCGCGGGTACTGCAAGCCTCCCA<br>AAACTGGGAGGCTTGCAGTACCCGC | IDT |
| Arid2-sgRNA1-O1<br>Arid2-sgRNA1-O2<br>Arid2-sgRNA2-O1<br>Arid2-sgRNA2-O2<br>Arid2-sgRNA3-O1<br>Arid2-sgRNA3-O2 | CACCGATAGCGAAGTCCACTTCATT<br>AAACAATGAAGTGGACTTCGCTATC<br>CACCGGTTTAAGAAGATCCCTGCGG<br>AAACCCGCAGGGATCTTCTTAAACC<br>CACCGCACTTTACTGCTCGCTAATG<br>AAACCATTAGCGAGCAGTAAAGTGC | IDT |
| hNT-sgRNA3-O1<br>hNT- sgRNA3-O2<br>hNT- sgRNA4-O1<br>hNT- sgRNA4-O2 | CACCGCAGGAGTCGCCGATACGCGT<br>AAACACGCGTATCGGCGACTCCTGC<br>CACCGTCGCGCTTGGGTTATACGCT<br>AAACAGCGTATAACCCAAGCGCGAC | IDT |

**Table 2. CRISPR/Cas9 targeting guide RNAs**

| Name | Sequence | Source |
| --- | --- | --- |
| Human ARID1A CRISPR Target Validation Primer-F | TACCAGGCCATCACAGCTTTTG | IDT |
| Human ARID1A CRISPR Target Validation Primer-R | TCCTCTCTGCCCCTATCACCTTTC | IDT |
| Human ARID1B CRISPR Target Validation Primer-F | GCAAGGCAGTTTCCCCGGCATG | IDT |
| Human ARID1B CRISPR Target Validation Primer-R | AGGAGGAGAGGGGAGAGAGAGG | IDT |
| Human ARID2 CRISPR Guide 1 Target Validation Primer-F | CAGTGTCATGTCACAAACTCTGC | IDT |
| Human ARID2 CRISPR Guide 1 Target Validation Primer-R | TGCCATTTACTTTTTACAAGACTGCC | IDT |
| Human ARID2 CRISPR Guide 3 Target Validation Primer-F | TGCTGTGCTGTTTTTCTGTTTCAG | IDT |
| Human ARID2 CRISPR Guide 3 Target Validation Primer-R | TCTCATGTGCAAAGGAAGGGAC | IDT |

**Table 3. Target validation primers**

| **Material Name** | **Catalog/Clone** | **Source** |
|---|---|---|
| TLCV2 | 87360 | Addgene |
| BsmBI-v2 | R0739S | New England Biolabs |
| Monarch DNA Gel Extraction Kit | T1020S | New England Biolabs |
| T4 Polynucleotide Kinase | M0201S | New England Biolabs |
| Quick Ligation Kit | M2200S | New England Biolabs |
| Stable Competent *E. coli* | C3040 | New England Biolabs |
| psPAX2 | 12260 | Addgene |
| pMD2.G | 12259 | Addgene |
| 10X RIPA buffer | 9806 | Cell Signaling Technology |
| Pierce Protease Inhibitor Mini Tablets | A32953 | ThermoFisher Scientific |
| DC Protein Assay Kit | 5000112 | Bio-Rad |
| 4x Laemmli Sample Buffer | 1610747 | Bio-Rad |
| Mini-Protean TGX Gels | 4561094 | Bio-Rad |
| **Antibodies for western blots** | | |
| IRDye® 800CW Goat anti-Rabbit IgG Secondary Antibody | 926-32211 | Li-Cor |
| IRDye® 680RD Goat anti-Mouse IgG Secondary Antibody | 926-68070 | Li-Cor |
| Ezh2 (D2C9) XP® Rabbit mAb | 5246 | Cell Signaling Technology |
| SMARCC1/BAF155 (D7F8S) Rabbit mAb | 11956 | Cell Signaling Technology |
| Brg1 (D1Q7F) Rabbit mAb | 49360 | Cell Signaling Technology |
| GAPDH (D16H11) XP® Rabbit mAb | 5174 | Cell Signaling Technology |
| c-Myc Antibody | 9402 | Cell Signaling Technology |
| Purified anti-c-Myc Antibody (9E10) | 626802 | BioLegend |
| ARID1A antibody | 12354 | Cell Signaling Technology |
| ARID1B antibody | 92964 | Cell Signaling Technology |

**Table 4. Materials used for CRISPR-Cas9 targeting and western blotting**

| Material Name | Catalog/Clone | Source |
|---|---|---|
| Rat anti-mouse CD34-Biotin (MEC14.7) | 119304 | BioLegend |
| Armenian hamster anti-mouse CD48-A700 (HM48-1) | 103426 | BioLegend |
| Rat anti-mouse CD150-PE-Cy7 (TC15-12F12.2) | 115913 | BioLegend |
| Rat anti-mouse CD150-PE (TC15-12F12.2) | 115904 | BioLegend |
| Rat anti-mouse c-Kit-APC-Cy7 (2B8) | 105826 | BioLegend |
| Rat anti-mouse FcγR-PE-Cy7 (93) | 101318 | BioLegend |
| Rat anti-mouse Flk2-Biotin (A2F10) | 13-1351-85 | eBioscience |
| Rat anti-mouse Ter-119-PE-Cy5 (TER-119) | 15-5921-83 | eBioscience |
| Streptavidin-BV605 | 405229 | BioLegend |
| Mouse anti-mouse CD45.1-PE (A20) | 12-0453-83 | eBioscience |
| Mouse anti-mouse CD45.1-PE-Cy7 (A20) | 25–0453-82 | eBioscience |
| Mouse anti-mouse CD45.2-FITC (104) | 11-0454-85 | eBioscience |
| Rat anti-mouse Sca-1-BV421 (D7) | 108128 | BioLegend |
| Rat anti-mouse CD11b-PE-Cy5 (M1/70) | 15-0112-83 | eBioscience |
| Rat anti-mouse Gr-1-PE-Cy5 (RB6-8C5) | 15-5931-83 | eBioscience |
| Rat anti-mouse CD4-PE-Cy5 (GK1.5) | 15-0041-83 | eBioscience |
| Rat anti-mouse CD5-PE-Cy5 (53-7.3) | 100610 | BioLegend |
| Rat anti-mouse B220-PE-Cy5 (RA3-6B2) | 15-0452-82 | eBioscience |
| Rat anti-mouse CD8a-PE-Cy5 (53-6.7) | 15-0081-83 | eBioscience |
| Armenian hamster anti-mouse CD3e-PE-Cy5 (145-2C11) | 15-0031-83 | eBioscience |

**Table 5. Antibodies used for multipotent progenitor (MPP) flow cytometry analysis and cell sorting**
